# A Biological Inventory of Prophages in *A. baumannii* Genomes Reveal Distinct Distributions in Classes, Length and Genomic Positions

**DOI:** 10.1101/2020.10.26.355222

**Authors:** Belinda Loh, Jiayuan Chen, Prasanth Manohar, Yunsong Yu, Xiaoting Hua, Sebastian Leptihn

## Abstract

*Acinetobacter baumannii* is of major clinical importance as the bacterial pathogen often causes hospital acquired infections, further complicated by the high prevalence of antibiotic resistant strains. Aside from natural tolerance to certain antibiotic classes, resistance is often acquired by the exchange of genetic information via conjugation but also by the high natural competence exhibited by *A. baumannii*. In addition, bacteriophages are able to introduce resistance genes but also toxins and virulence factors via phage mediated transduction. In this work, we analysed the complete genomes of 177 *A. baumannii* strains for the occurrence of prophages, and analysed their taxonomy, size and positions of insertion. Among all the prophages that were detected, Siphoviridae and Myoviridae were the two most commonly found families, while the average genome size was determined as 3.98 Mbp. Our data shows the wide variation in the number of prophages in *A. baumannii* genomes and the prevalence of certain prophages within strains that are most “successful” or potentially beneficial to the host. Our study also revealed that only two specific sites of insertion within the genome of the host bacterium are being used, with few exceptions only. Lastly, we analysed the existence of genes that are encoded in the prophages, which confer antimicrobial resistance (AMR). Several phages carry AMR genes, including OXA-23 and NDM-1, illustrating the importance of lysogenic phages in the acquisition of resistance genes.

## Introduction

The opportunistic pathogen *Acinetobacter baumannii* is the causative agent for bloodstream infections, meningitis and urinary tract infections, and is responsible for 2-10% of all Gram-negative hospital-acquired infections (Joly-Guillou 2005). Such infections include ventilator-associated pneumonia and bacteremia with a mortality rate of 35-52% (Dijkshoorn et al. 2007; Kempf et al. 2013; Antunes et al. 2014). As a multitude of strains cause nosocomial infections, *A. baumannii* has become of major important pathogen in hospital care and is of global concern. Many clinical isolates have acquired genes coding for virulence factors, such as toxins or efflux pumps, through various genetic uptake mechanisms (Morris et al. 2019). While *A. baumannii* easily acquires genetic material by conjugation, natural transformation is also widespread as many strains are highly naturally competent (Hu et al. 2019). Such mechanisms ultimately give rise to an increasing number of strains that display high levels of antimicrobial resistance (AMR), against which antibiotics show little or no effect. Genetic information for AMR genes is often embedded in genetic elements such as transposons or plasmids (Partridge et al. 2018; Rozwandowicz et al. 2018). In addition, bacteriophages (or phages) are able to transfer non-viral genetic information through a process called transduction, which can include genes coding for toxins or antimicrobial resistance (Wagner and Waldor 2002; Derbise et al. 2007; Wachino et al. 2019). Therefore, phages play an important role in the development of AMR.

Regardless of their morphology or infection mechanism, phages can be divided into two types based on their life cycle: Lytic phages and lysogenic phages (sometimes also called temperate). Both eventually kill the host cell by lysis, employing various enzymes that create holes in the membrane and disintegrate the bacterial cell envelope, to allow the release of phage progeny. Few exceptions exist, such as the filamentous phages that are assembled in the membrane and secreted from the host while the bacterium continues to grow and divide (Loh et al. 2017; Loh et al. 2019; Kuhn and Leptihn 2019). Nonetheless, phages that destroy the host by lysis upon completion of their life cycle, either start their viral replication immediately after entry (lytic phages) or integrate their genome into that of the host first (lysogenic). Lysogenic phages can remain “dormant” without replicating their genome or initiating phage coat protein synthesis. This way, lysogenic phages are being inherited by daughter cells, and might only replicate to form phage particles after many generations. The trigger for phage replication and synthesis is usually a stress signal produced by the host, such as a SOS response after DNA damage (Howard-Varona et al. 2017). However, if the host resides under favourable conditions, lysogenic phages continue their passive co-existence as so-called prophages embedded inside the DNA of the host.

Prophages are a major source of new genes for bacteria, occupying up to 20% of bacterial chromosomes and therefore may provide new functions to its host (Brüssow et al. 2004; Brüssow 2007; Cortez et al. 2009, Fortier and Sekulovic 2013; Wang and Wood 2016). These functions include virulence factors and drug resistant mechanisms which include extracellular toxins and effector proteins involved in adhesion factors, enzymes, super antigens and invasion (Fortier 2017; Tinsley et al. 2006, Wang and Wood 2016; Argov et al. 2017). In some cases, the acquisition of virulence genes allows non-virulent bacteria to become a virulent pathogen. The most prominent example is that of the CTXΦ cholera toxin, encoded by a filamentous phage, making *Vibrio cholerae* the clinical pathogen that poses a substantial socio-economic burden on developing countries with poor hygiene due to frequent cholera outbreaks (Davis and Waldor 2003). Another example is the Shiga toxin-encoding prophages found in highly virulent *Escherichia coli* strains, causing food-borne infections across the world (Gamage et al. 2004; Tozzoli et al. 2014). As part of the bacteriophage life cycle, prophages of lytic phages are a double edged sword; while they provide the advantage of increasing chances of survival in challenging environments, they could also lead to the killing of the host through the release of progeny at the end of the phage life cycle.

Our relationship with microorganisms is a complex and vital one, from the important role of gut microbiota to the increase in mortality due to virulent microorganisms, understanding bacterial genomes is crucial. Yet in order to understand them in detail, we also need to be able to identify viral genes and to understand the impact of these genes on its host. As part of the bacterial genome, prophages are subjected to the general effects of mutation, recombination and deletion events. For some phages it has been clearly established that prophage genes have an influence on the host, such as motility or biofilm formation, both important aspects with regards to virulence. However for many other prophages, it is less well understood. Previous work on *A. baumannii* prophages have identified putative virulence factors and antibiotic resistance genes in host genomes deposited on GenBank (Costa et al. 2018, López-Leal et al. 2020). The work presented here analyses clinical *A. baumannii* strain genomes in search of possible prophages. We describe the identification of active prophages, analysed their taxonomy, size and detail the regions in the bacterial genomes where these prophages have been found. Our data reveal a wide variation of the number of prophages in *A. baumannii* genomes and the prevalence of certain prophages within strains. From an evolutionary perspective, these might represent the “most successful” phages, or the ones that bring a benefit to the host. Our data analysis also allowed us to identify two major sites of insertion within the genome of the host bacterium, with most phage genomes inserting in these two regions, while only a few exceptions are being observed. In addition, our study indicates two distinct genome size distributions of prophages, as we observe a bimodal distribution when analysing all prophage genomes. Furthermore, we describe genes coding for virulence factors, in particular for antimicrobial resistance in the prophages.

## Materials and Methods

### *A. baumannii* genomes

Complete genome sequences of *A. baumannii* only were selected for this study. Detailed information of each *A. baumannii* strain used in this study is disclosed in supplementary material (Supplemental Table S1). For the characterisation of genome lengths of *A. baumannii*, the following values were obtained: mean, median, mode, the smallest and the largest genome. The distribution of genome lengths was plotted using the geom_density function provided in the ggplot2 package in R (Wickham 2016).

### Alignment of *A. baumannii* genomes

All strains were aligned so that their starting position is identical, with the gene dnaA defined as the start. BLAST Scoring Parameters provided by NCBI (Gertz et al. 2005; available at https://blast.ncbi.nlm.nih.gov/Blast.cgi) was used to blast the locations of dnaA and the adjustment of genome sequences was achieved by SnapGene software (from Insightful Science; available at snapgene.com).

### Identification of Prophage Genes

The tool used to identify prophages in *A. baumannii* genomes was Prophage Hunter (Song et al. 2019; available at https://pro-hunter.bgi.com/). Here, we obtained data on the start, end, length, score, category, and the name of the closest phage. Phaster (Arndt et al. 2016; available at http://phaster.ca/) was used to further confirm some conflicting results. According to the algorithm created by the authors of Prophage Hunter, an “active” prophage is defined by a score close to 1, while the probability decreases the lower the score gets. This means that an active prophage region received a scoring of higher that 0.8 while 0.5 to 0.8 is defined as “ambiguous”, and a score lower than 0.5 as “inactive”.

### Prophage number analysis

Calculations of the mean, median, and mode on the prophage number (total, only active, and only ambiguous) were performed after removing the overlaps of the same prophage in the same strain. The ten strains with the fewest and with the largest numbers of total, only active, and only ambiguous prophages were selected to show the two extremes, while the density plot achieved through geom_density function provided by ggplot2 package in R were conducted to describe the general distribution. The boxplot produced by R reflected the relationship between total, only active, and only ambiguous prophage number and *A. baumannii* genome length.

### Phylogenetic analysis of host strains

Prokka v1.13 (Seemann 2014) was used to generate the gff files for the genome sequences of 177 *A. baumannii* strains. The core genome alignment was constructed with Roary v3.12.0 (Page et al. 2015). A maximum-likelihood phylogenetic tree was created using FastTree v2.1.10 (Price et al. 2010). The tree was annotated and visualized with ggtree.

### Prophage classification and phylogeny analysis

Prophage classification presented was provided by the program Prophage Hunter which was based on the NCBI’s database. Different orders and families were taken into consideration. The prophage number in different families and their proportions were revealed by histograms and pie charts generated in Microsoft Excel respectively. Different bacterial hosts were referred to in the calculation of the number of prophages and the number of phage species they had. Based on the species of prophages, ten most common ones for total, only active, and only ambiguous were selected, with a heatmap which was completed through geom_tile provided by ggplot2 package in R showing their number with the activity value in different strains. The phylogeny of phages was mapped according to their sequences. The alignment of the phage sequences were performed using Multiple Alignment using Fast Fourier Transform (MAFFT) (Katoh and Standley 2013) with default options. Maximum-likelihood phylogenetic trees were created using FastTree v2.1.10. The tree was annotated and visualised using ggtree.

### Prophage location analysis

The positions of all prophages were first showed in a stacked bar chart in Microsoft Excel. Considering the overlaps of different prophages in the same strain, all the prophage starts and ends were mapped in a density plot created by ggplot2 geom_density and aes functions in R. The stacked bar charts of different prophage species were used to estimate their preference of insertion which were then summarized in tables.

### Prophage length analysis

The use ggplot2 geom_density function in R facilitated the creation of density plots of prophage length. With the help of aes function, the mapping of prophage categories (active and ambiguous) and families were achieved in the density plots.

### Identification of virulence factors and antibiotic resistance genes

No program is currently available that allows the search of virulence genes which are embedded in prophage sequences within bacterial genomes. Thus, we identified prophages first followed by manually correlating genomic positions of virulence genes with those that were also identified to belong to prophage genes.

### Mapping of prophage-encoded antimicrobial resistance genes (ARG)

To search for the specific virulence genes we identified (above), we first downloaded all available 4128 *A. baumannii* Illumina sequencing reads from the Sequence Read Archive (SRA) with the cut-off date for deposited sequences on 2019/11/17. The raw Illumina sequencing reads were mapped against the ARG prophage sequences employing BWA-MEM v0.7.17 (80% coverage cutoff) (Li 2013).

## Results

### In silico discovery of prophages in 177 A. baumannii genomes identifies 1156 prophage sequences

Our first aim was to analyse how frequently prophages occur in the genomes of *A. baumannii* strains. We randomly chose 177 genome sequences of *A. baumannii* strains, many of them clinical isolates. For the subsequent analyses, we aligned all sequences so that their starting positions are identical. To this end, we defined the gene dnaA as the start, which encodes for a replication initiation factor that facilitates DNA replication in bacteria. From the sequence alignments, we observed a large variation in genome sizes. The average length of the genomes was 3,981,579 bp with a median value of 4,001,318 bp; the smallest genome had a length of 3,072,399 bp (22.9% shorter than average), and the largest genome displayed a size of 4,389,990 bp (10.25% larger than average) (Figure 1A).

**Figure 1:**
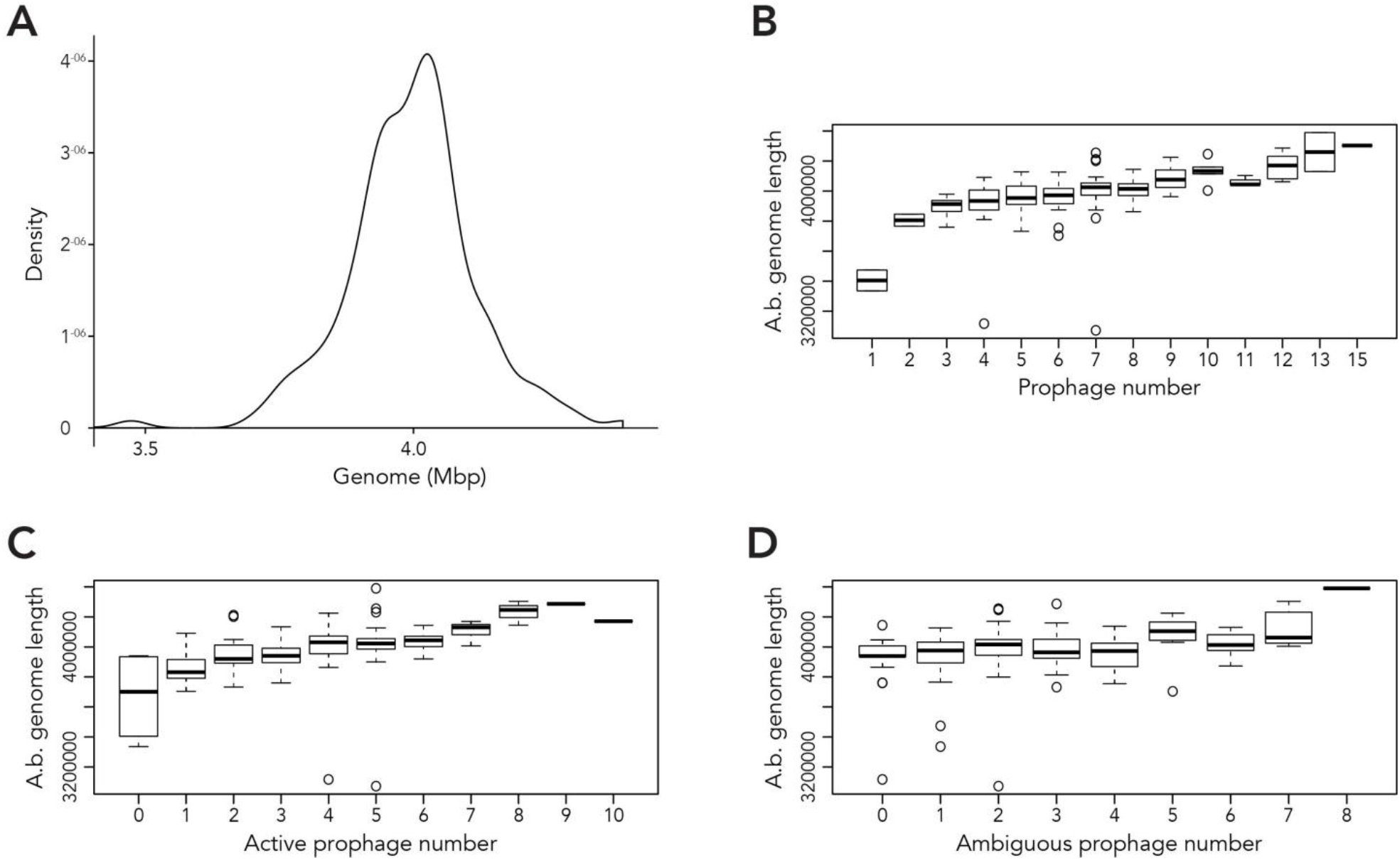
Length of *A. baumannii* genomes and its correlation with number of prophages present. A) Density graph of 177 genome sequences of *A. baumannii* strains, indicating the distribution of the lengths of genome sequences analysed. B - D) Correlation between *A. baumannii* genome length and number of B) all prophages, C) active prophages and D) ambiguous prophages identified in genomes.

Next, we used the online platform Prophage Hunter (Song et al. 2019) to identify active and ambiguous prophage genomes. The algorithm provides an output value for each prophage identified, which allows the researcher to establish whether a sequence contains an “active” or an ambiguous prophage. While “active” prophages exhibit the complete genomic sequence of a prophage, and are therefore likely to allow the production of phage particles, ambiguous prophage sequences are truncated, mutated or otherwise incomplete, and unlikely to be able to form infectious phages. Among the 177 *A. baumannii* genomes analysed, we identified 1156 prophages, with 459 of them being defined as “ambiguous” while the remaining 697 sequences were labelled as “active” according to the program. To determine the prevalence of prophages in the *A. baumannii* genomes, we analysed the number of prophages per genome. Using a heat map to illustrate our results, we found that while some prophages are rarely found, others are quite common in the genomes of *A. baumannii* isolates (Figure 2). One phage that was only found once, for example, is a prophage with high sequence similarity to the Yersinia Podovirus fHe-Yen3-01, while the *Acinetobacter* phage Bphi-B1251, a Siphovirus, has been found in 79.1% (140 in 177) of all analysed *A. baumannii* genomes. Such high prevalence observed by prophages such as Bphi-B1251 could indicate high infectivity and wide host range of the active phage particle. Table 1 shows ten strains with the fewest and with the largest numbers of prophages identified (Table 1). We found that the average prophage number in an *A. baumannii* genome is 6.53, with some bacterial genomes containing only 1 (n = 2) prophage sequence, such as in case of the *A. baumannii* strains DS002 and VB1190. In contrast to these, other strains have been found to contain as many as 10 prophage sequences, such as in strain 9201 (n = 1), that were labelled “active” by Prophage Hunter; additionally this strain contains 2 prophage sequences that were defined as ambiguous. The highest prophage number was found in the strain AF-401 which contains 15 prophages, however only 8 were defined as active. Our results show that prophage sequences are relatively common and that most *A. baumannii* strains show a median of 7 and a mode of 8 prophages per genome. Additional genome analysis of the clinical isolates illustrates a possible relationship between prophages and host strains (Supplemental Figure S1). Prophages such as *Bacillus* phage PfEFR 4 and *Enterobacteria* phage CUS 3 were observed more frequently in strains whose genomes are in the same clade or are closely related, indicating a narrow host range.

**Table 1:** *A. baumannii* stains with the highest and the fewest number of prophages identified.

| Highest number of prophages identified |  | Fewest number of prophages identified |  |
| --- | --- | --- | --- |
| Strain | Prophage number | Strain | Prophage number |
| AbPK1 | 11 | DS002 | 1 |
| AR_0056 | 11 | VB1190 | 1 |
| 9201 | 12 | CA-17 | 2 |
| 10042 | 12 | E47 | 2 |
| AR_0101 | 12 | 11A14CRGN003 | 3 |
| DU202 | 12 | 11A1213CRGN008 | 3 |
| VB35435 | 12 | 11A1213CRGN055 | 3 |
| 11W359501 | 13 | 11A1213CRGN064 | 3 |
| AB030 | 13 | 11A1314CRGN088 | 3 |
| AF-401 | 15 | 11A1314CRGN089 | 3 |

**Figure 2:**
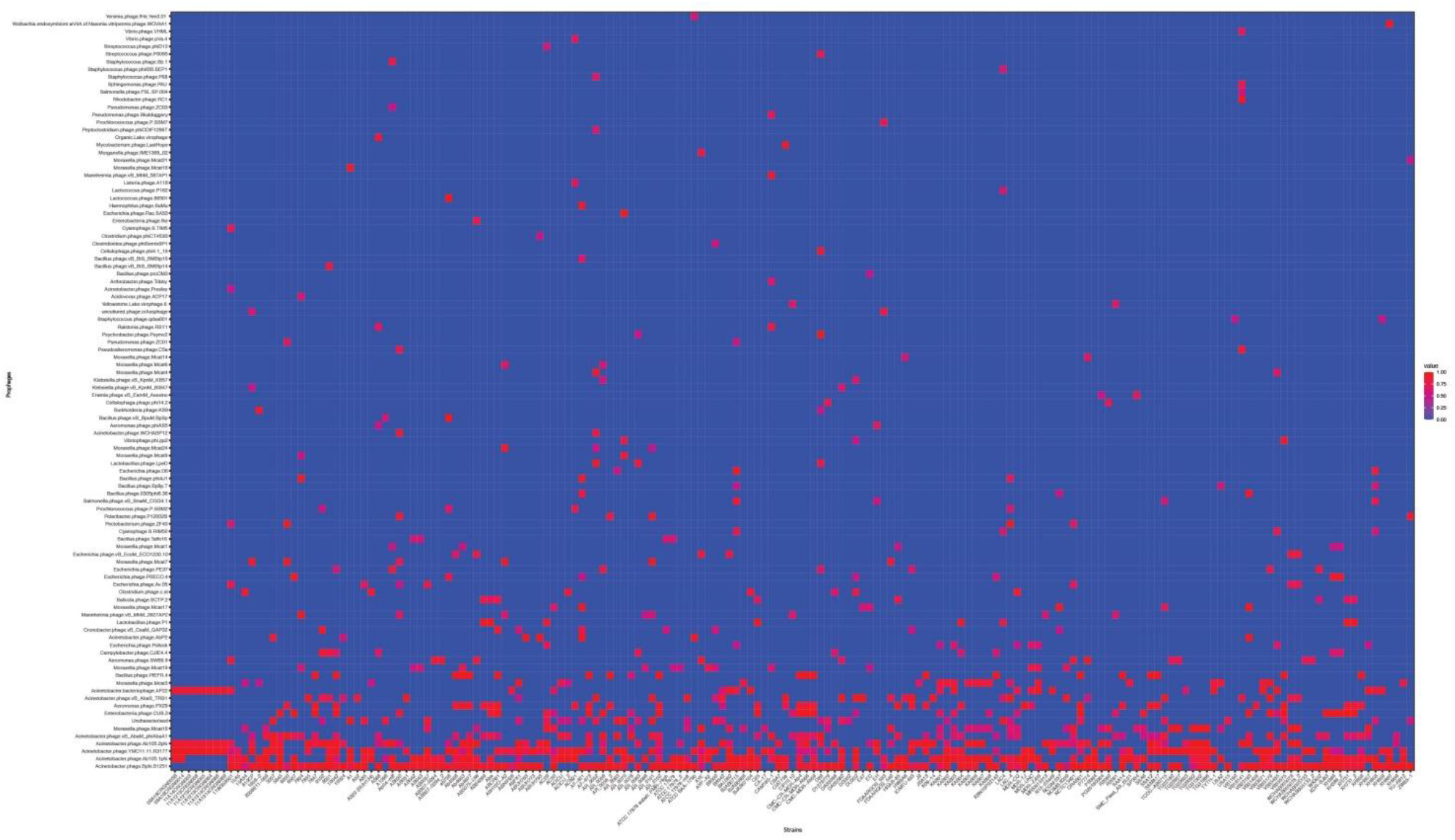
The prevalence of prophages analyzed. Heat map of prophages found in all A. baumannii strains analysed. Prophages (y-axis) are plotted against each A. baumannii strain (x-axis). Red squares indicate the presence of the indicated prophage. Blue squares indicate the lack thereof. Please refer to the PDF of the figure and use the zoom function to identify names of strains and phages.

We next correlated host genome length with number of prophages identified to determine if there is a relationship between the two variables. Perhaps unsurprisingly, the length of *A. baumannii* genomes increased as more prophages are identified, disregarding whether the prophage genomes are “active” or “ambiguous” (Figure 1B). However, when prophages are classified, the correlation between host genome length and number of prophage genomes identified show less distinction (Figure 1C & D).

### Siphoviridae and Myoviridae are the two most commonly found classes of prophages in A. baumannii genomes

We analysed the relationship of all prophages we identified and created a phylogenetic tree (Supplemental Figure S2). Phage phylogeny is very complex. While bacteria share many common genes, microbial viruses are less related to each other creating large phylogenetic distances. The phylogenetic analysis shows that in some instances the phylogenetic clustering does not necessarily result in grouping of the phages according to their classes. This is however not surprising as phages often display diversity by “mosaicity of their genomes” (Dion et al. 2020). After identifying all prophage sequences in the genomes of the 177 *A. baumannii* isolates, we set out to analyse the most prevalent classes of phages present. Our analysis of all prophage sequences (n = 1156) revealed that the majority of them, ~57% (n = 660) of the prophages, belong to the Siphoviridae group (Figure 2A). The Siphoviridae is a class of head-and-tail phages, with the best known representative being phage lambda, that exhibit long, non-contractile but comparably flexible tail structures (Nobrega et al. 2018). The second most commonly found phages are Myoviridae, with a percentage of ~33% (n = 385) (Figure 2A). With the best-known Myovirus, the *E. coli* phage T4, these phages have a stiff, contractible tail that allows the active penetration of the bacterial host envelope (Hu et al. 2015). Together, these two phage classes make up 90% of all prophage genomes. The third most common class, albeit only 4.7% (n=55) of all prophage genomes, belongs to Podoviridae. The best known Podovirus is probably T7, which has a short, stubby tail and internal core proteins that get ejected for the formation of a DNA-translocating channel across the bacterial cell envelope (Guo et al. 2014; Lupo D et al. 2015; Leptihn et al. 2016). Prophages that could not be conclusively classified to a viral group accounted for 3.3% (n=38). Siphoviridae, Myoviridae and Podoviridae all belong to the order Caudovirales, phages that exhibit a head-and-tail structure. Within the two phage classes we found several phages that were most successful, i.e. most common. Examples are the *A. baumannii* phages Bphi-B1251 and YMC11/11/R3177, which both belong to the Siphoviridae (Table 2A). The most common Myovirus was Ab105-1phi. Not only are these the most common prophages found, they are also the most common active prophages identified (Table 2B). In addition, the distribution of the classes was similar if only active prophages were analysed. Here, 62% (432/697) belonged to the Siphoviridae and 32% (223/697) to the Myoviridae (Figure 2B).

**Table 2:** The most common prophages identified in *A. baumannii* stains.

| A 10 the most common prophages (total) |  | B 10 the most common prophages (active) |  |
| --- | --- | --- | --- |
| Phages | Number found | Phages | Number found |
| Acinetobacter phage Bphi-B1251 | 228 | Acinetobacter phage Bphi-B1251 | 177 |
| Acinetobacter phage Ab105-1phi | 143 | Acinetobacter phage Ab105-1phi | 111 |
| Acinetobacter phage YMC11/11/R3177 | 118 | Acinetobacter phage YMC11/11/R3177 | 78 |
| Acinetobacter phage Ab105-2phi | 95 | Acinetobacter phage Ab105-2phi | 65 |
| Acinetobacter phage vB_AbaM_phiAbaA1 | 56 | Aeromonas phage PX29 | 31 |
| Moraxella phage Mcat16 | 42 | Enterobacteria phage CUS-3 | 25 |
| Uncharacterised | 38 | Acinetobacter bacteriophage AP22 | 23 |
| Enterobacteria phage CUS-3 | 35 | Bacillus phage PfEFR-4 | 20 |
| Aeromonas phage PX29 | 34 | Acinetobacter phage vB_AbaS_TRS1 | 19 |
| Acinetobacter phage vB_AbaS_TRS1 | 33 | Moraxella phage Mcat3 | 16 |

### The genomic position of prophages shows two main locations for genome integration

To determine where all prophages -regardless of their class- are found in the bacterial genome, or if they are possibly distributed at random within the host DNA, we visualised the position of the prophages in all 177 genomes (Figure 4A). We then plotted all prophage positions found in all genomes against the position in the bacterial host genome sequence. Surprisingly, we observed a bimodal distribution, with two clear peaks in the position of prophages (Figure 4B), indicating that there are two main sites of attachment for prophages and their genomic insertion. While this reflects the situation for all prophages, we then analysed the position of several individual prophages within the bacterial genome. First, we assessed one of the most commonly found phages YMC11/11/R3177. The position of this phage reflects the overall distribution of all phages in the analysed genomes, with two main areas of insertion. However, in some cases the position is outside the main area of insertion, possibly due to recombination of the bacterial genome (Figure 5A). The second phage we analysed was phage vB_AbauS_TRS1. Here, the distribution of the phage within the bacterial genome seems to be more random as compared to the overall distribution (Figure 5B). In case of the *Aeromonas* phage PX29, insertion seems to be very “strict”, i.e. only observed in one location within the genome (Figure 5C). The observations made when analysing the prophage positions show that the insertion of phages could be described as “directed”, and less random, indicating that attachment sites, if they are required, are found more commonly in certain positions of the bacterial genome.

**Figure 3:**
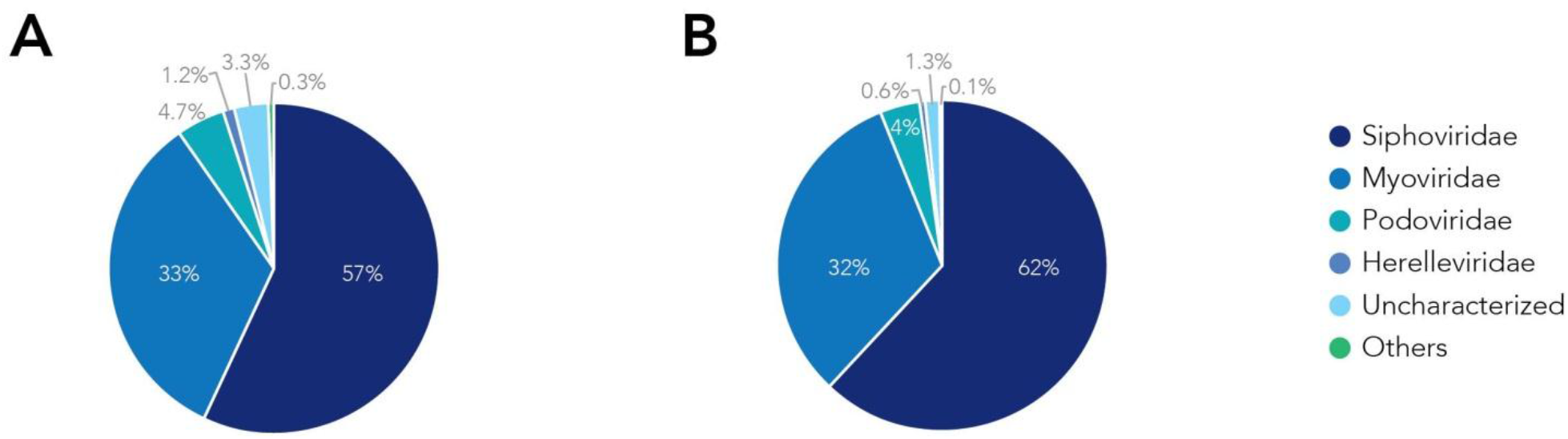
The families of prophages found. Pie charts of prophages identified showing the percentage make up of each family. A) Classification for all prophages. B) Classification of active prophages only.

**Figure 4:**
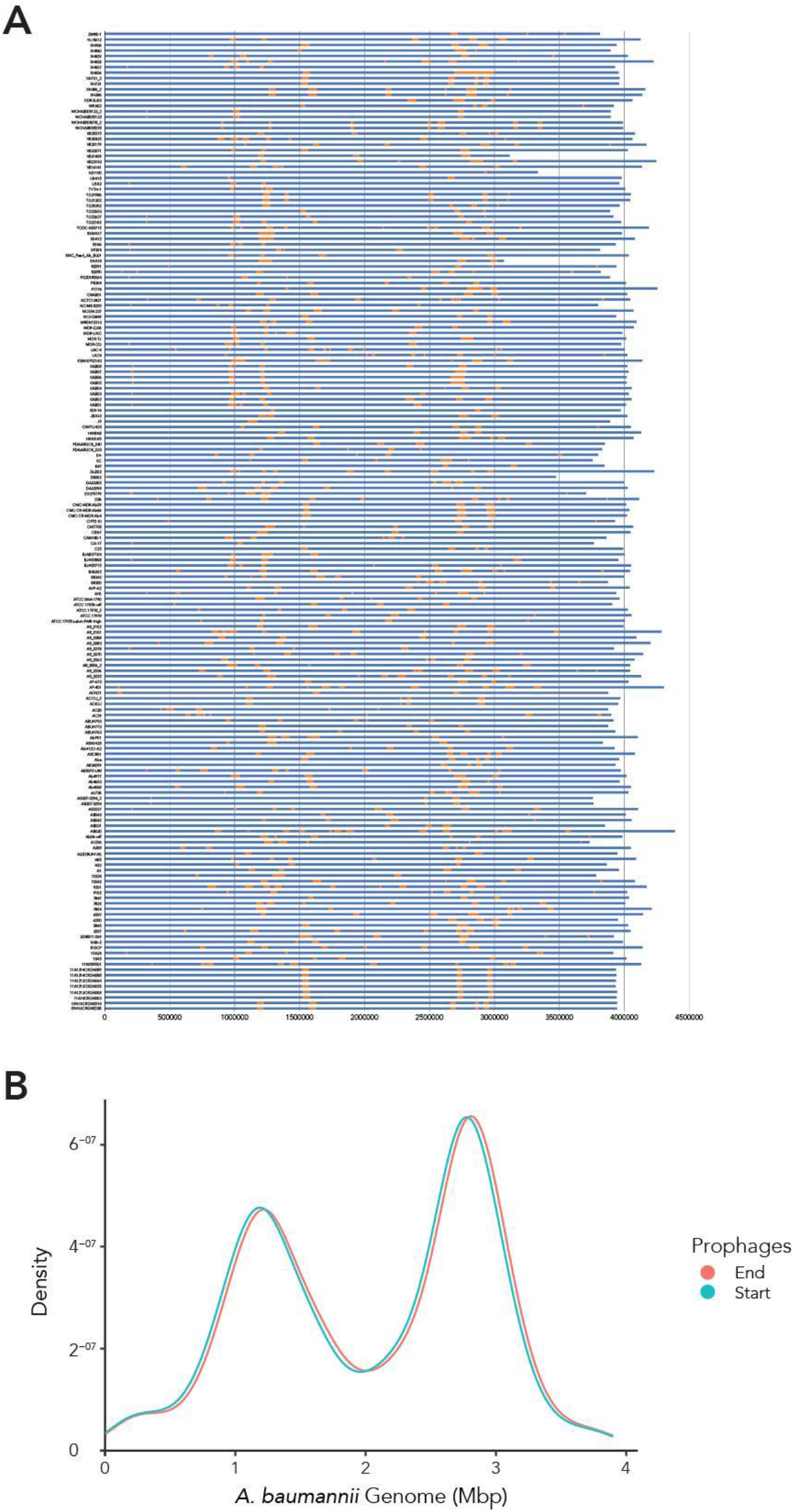
The location of prophages found in each *A. baumannii* genome. A) Heat map of each bacterial strain (y-axis). Yellow segments indicate prophage sequences identified on the genome (in blue). B) Density graph compiled from the heat map data. Please refer to the PDF of the figure and use the zoom function to identify names of strains and phages.

**Figure 5:**
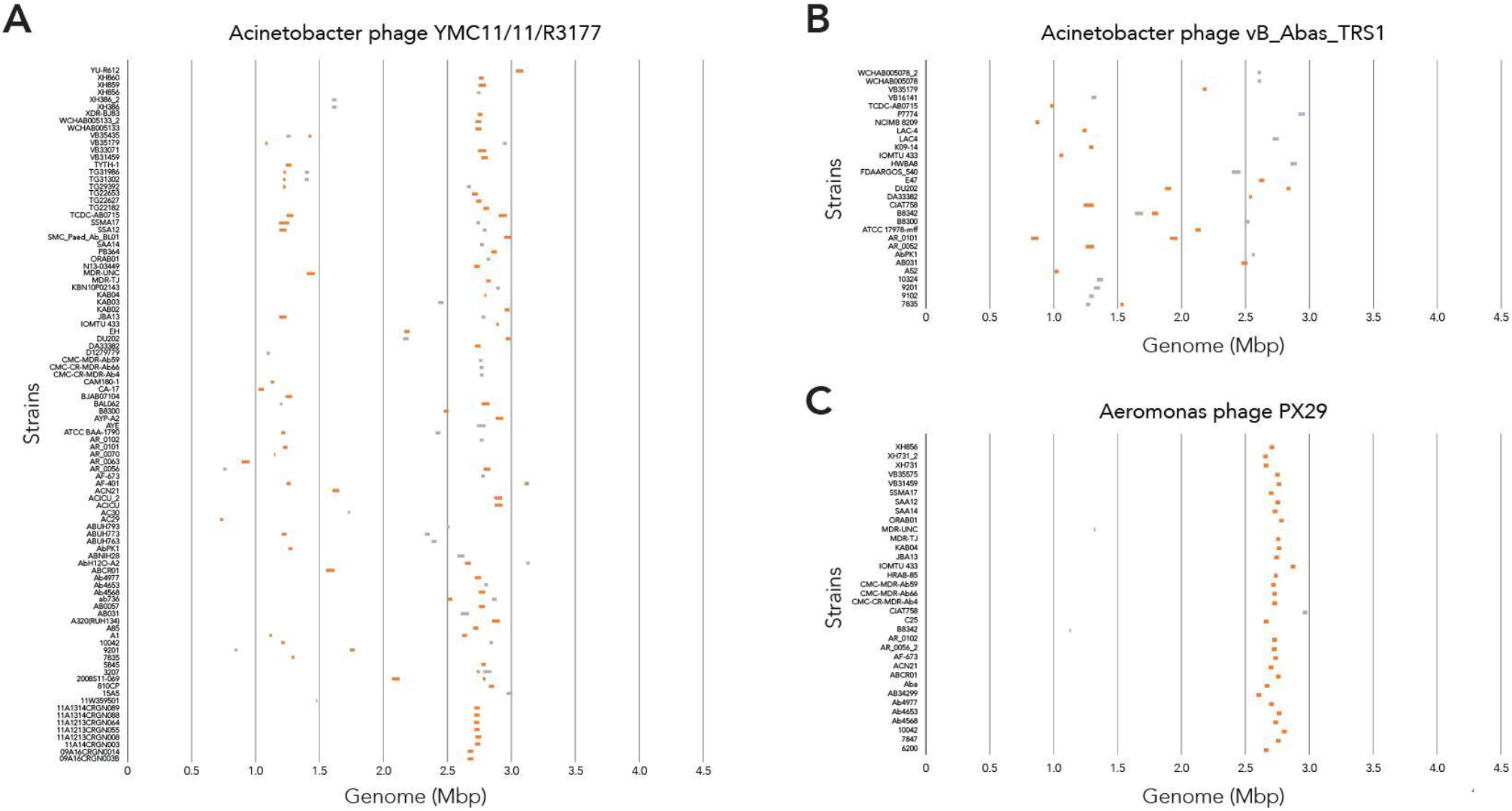
Location of prophage genome insertion differs between phages. A) Comparison of phage location for Acinetobacter phage YMC11/11/R3177. B) Comparison of insertion locations for Acinetobacter phage vB_Abas_TRS1. C) Comparison of prophage insertion sites for Aeromonas phage PX29. Boxes in orange indicate active prophages identified. Grey boxes indicate ambiguous prophage sequences. Please refer to the PDF of the figure and use the zoom function to identify names of strains and phages.

### The sequence length of prophages reveals distinct groups

Using the data provided by the program Prophage Hunter, we were interested in evaluating the size distribution of prophage genomes. Therefore, we plotted sizes against the frequency of prophages present in the bacterial genomes and calculated average prophage genome sizes. In the case of ambiguous prophages, a main population at 15 kb became visible followed by a minor peak of substantial size at approximately 60 kb, leaving the average and median length of ambiguous prophage genomes at 29.2 kb and 25.8 kb respectively (Figure 6A and Table 3). In contrast to this, two main peaks were observed when analysing only active prophages. Here, one peak is observed at around 17 kb while the other at around 36 kb was observed (Figure 6A). The average genome length of active prophages is 34 kb (Table 3). As these peaks include all phage categories, we re-analysed the genomic length of the active prophages according to their classes: Siphoviridae, Myoviridae and Podoviridae, which together constitutes almost 95% of all prophages (see Figure 2A). When analysing the length of all Siphoviridae sequences, we observed two main populations, one sharp peak at around 20 kb and one broad peak with a shoulder containing larger sequences from 18 to 56 kb (Figure 6B). The average prophage length of Siphoviridae is 36.7 kb (Table 3). Myoviridae sequences similarly exhibited two sharp peaks (17 and 36 kb), with a third minor one of around 60 kb (Figure 6C). The average prophage genome size of active Myoviridae is 32.4 kb (Table 3). The Podoviridae showed several minor peaks with a large sharp peak at about 12 kb and the average prophage genome length is calculated to be 17.4 kb (Figure 6D and Table 3).The results of these analyses show that there are distinct distributions of bacteriophage genome sizes. Two clearly separated groups of prophages can be observed just based on size, in the case of Myoviridae. In the case of Siphoviridae, we saw a less defined area with possibly multiple species within the broad distribution.

**Table 3:** Prophage genome lengths for active and ambiguous prophages, among the 3 major families in the order of Caudovirales.

|  |  | All Prophages | Active Prophages | Ambiguous Prophages |
| --- | --- | --- | --- | --- |
| All Prophages | Average | 32211.62 | 34182,66 | 29218,56 |
|  | Median | 31296 | 35074 | 25759 |
| Siphoviridae | Average | 34586.75 | 36668,1 | 30625,76 |
|  | Median | 33018 | 36562,5 | 27679 |
| Myoviridae | Average | 32148,62 | 32351,98 | 31870,4 |
|  | Median | 34249 | 35075 | 30792 |
| Podoviridae | Average | 19422,25 | 17357,86 | 21563,11 |
|  | Median | 15498 | 15498 | 14963 |

**Figure 6:**
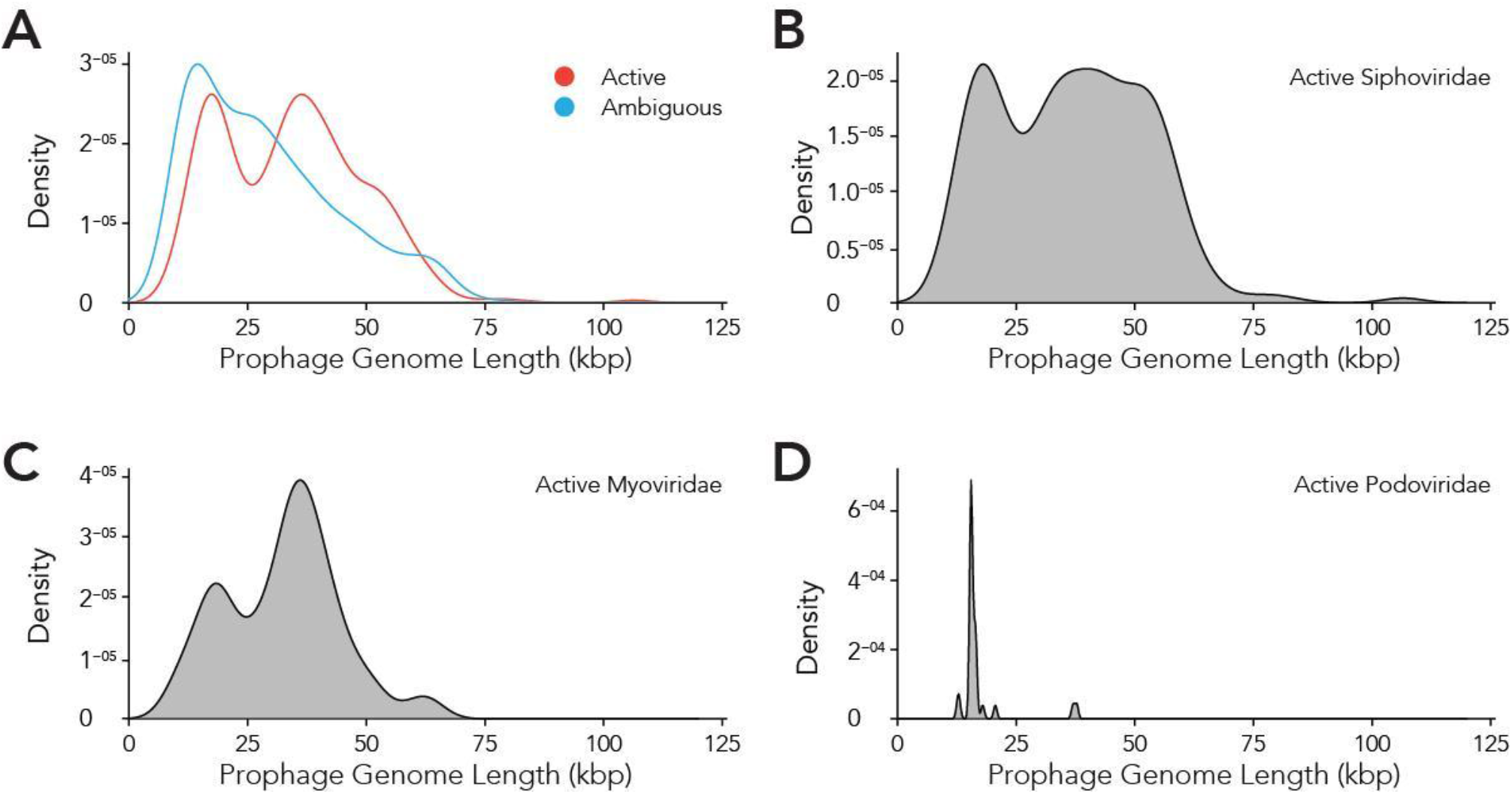
Active prophages categorised by prophage lengths. A) Comparison of prophage lengths between active and ambiguous prophages. B - D) Distribution of prophage length for active Siphoviridae (B), Myoviridae (C) and Podoviridae (D).

### Prophage encoded antibiotic resistance genes

As prophages are able to encode genes that might allow its host to become more virulent and therefore more evolutionary successful, we aimed to analyse prophage-encoded virulence factors. However, in contrast to e.g. *E. coli*, a databank for *A. baumannii* virulence factors currently does not exist. We therefore searched for prophage sequences that contain genes that contribute to antibiotic resistance. Table 4 lists the start and end of the genes that are encoded within a respective prophage. Among others, we found AMR genes for OXA-23 and NDM-1. OXA-23 is the most widespread carbapenem resistance gene globally (Hamidian and Nigro 2019). NDM-1 encodes a carbapenemase, a beta-lactamase enzyme with a broad substrate specificity capable of hydrolyzing penicillins, carbapenems, cephalosporins, and monobactams. Other beta-lactamase genes were *bla*_ADC-5_, *bla*_OXA-67_, *bla*_OXA-115_, and *bla*_TEM-12_. In addition, we were able to identify genes coding for N-Acetyltransferases(aac(3)-I, *aac(3)-Id*, *aacA16)*, Aminoglycoside phosphotransferases (*aph(3’)-Ia*, *aph(3’)-VI*, *aph(6)-Id*, *aph(3”)-Ib*), both groups mediating aminoglycoside resistance. Other genes that contribute to antibiotic resistance were sulfonamide resistance gene (*sul2*), and the macrolide-resistance conferring genes *msr(E)*, encoding an efflux pump, and *mph(E)*, coding for a macrolide-inactivating phosphotransferase.

**Table 4:**
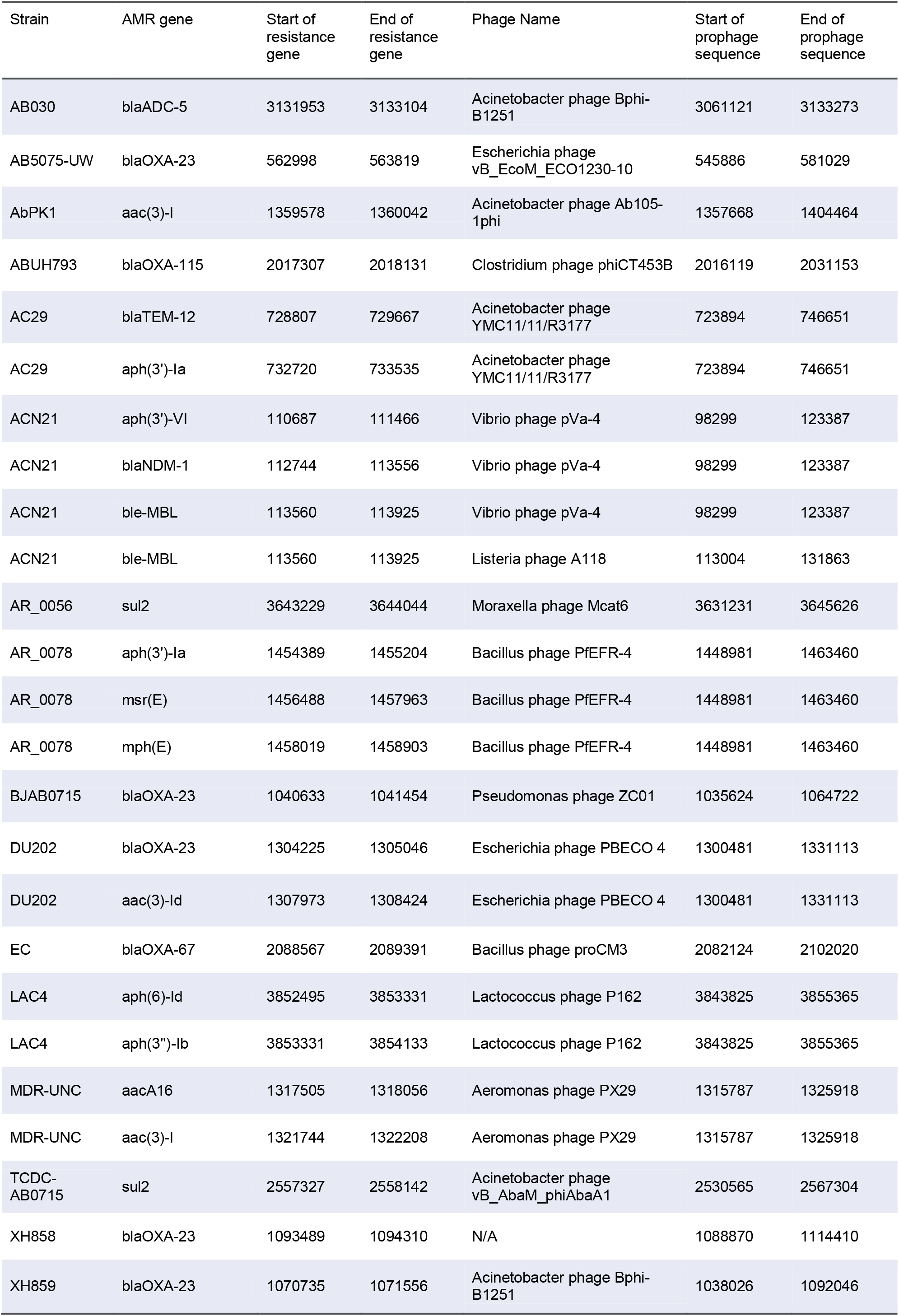
Antimicrobial resistance genes found in prophages embedded in *A. baumannii* stains.

Interestingly, one *A. baumannii* strain, ACN21, contains a prophage which encodes three antibiotic resistance genes (*aph(3’)-VI*, *bla*_NDM-1_, ble-MBL). The phage is most closely related to Vibrio phage pVa-4, a Myovirus that infects *V. alginolyticus* (Kim et al. 2019). In contrast to its relative, phage pVa-4 is a lytic phage was grouped to be part of the phiKZ-like phages (Phikzviruses), which are considered as "jumbo" phages. A second *A. baumannii* strain, AR_0078, contains a prophage sequence that shares a high degree of similarity to the Bacillus phage PfEFR-4, a Siphovirus with a prolate head, in contrast to its *E. coli* relative lambda (Geng et al. 2017). The prophage encodes three antimicrobial resistance genes that confer macrolide-resistance, *msr(E)*, encoding an efflux pump, and *mph(E)*, coding for a macrolide-inactivating phosphotransferase. The third gene encodes the Aminoglycoside phosphotransferases (*aph(3’)-Ia*), inactivating aminoglycoside antibiotics.

*A. baumannii* strain DU202 contains a prophage sequence that is related to the lytic *E. coli* myovirus PBCO 4 (Kim et al. 2013). Within the genome sequence, two antimicrobial resistance genes are encoded: *aac(3)-Id* codes for an N-Acetyltransferase mediating aminoglycoside resistance and OXA-23, which is encodes the most widespread resistance mechanism towards the β-lactamase inhibitor sulbactam. The gene was also embedded in the prophage sequences of two phages in the *A. baumannii* strain XH859; here, the *A. baumannii* phage Bphi-B1251 was found to be the most closely related phage, a lytic Podovirus, that was previously shown to be able to infect and lyse an OXA-23-harbouring *A. baumannii* isolate from a septic patient (Jeon et al. 2012).

To determine how prevalent the prophage encoded antimicrobial resistance genes are in other genomes, we mapped all available 4128 *A. baumannii* Illumina sequence reads that were accessible by 2019/11/17 on the Sequence Read Archive (SRA) to the AMR prophage sequences using an 80% cutoff for the coverage. We identified 174 *A. baumannii* genomes that contain prophage sequences, or about 4.2% of all available reads, not including the genomes we used for the initial analysis. Supplemental Figure S3 illustrates the prevalence of the prophage-encoded AMR genes (ARGs) and their respective prophages. Fairly “successful”, i. e. widely distributed, were two prophages: ABUH793 and AR_0056. ABUH793 is a close relative of the Clostridium phage phiCT453B, containing the resistance gene blaOXA-115. The second prophage sequence that was found often in *A. baumannii* genomes in comparison to other prophages encoding ARGs, was AR_0056, a relative of the Moraxella phage Mcat6, encoding the ARG sul2. While the prophage does not necessarily render the host antibiotic resistant as genetic regulators might be missing, prophages containing ARGs can present an evolutionary advantage for the host (Wendling et al. 2020).

Our finding demonstrates the importance of phages in the acquisition of antimicrobial resistance; the above described genes may confer the ability to grow in the presence of antibiotics when the bacterial host is infected by a phage that encodes not only the information for its own replication but also genes that inactivate or remove antibiotic compounds.

## Discussion

Our search for prophages in the genomes of *A. baumannii* strains revealed several interesting findings. One surprising observation was the positions of the prophages within the genome of the bacterial host. When analysing the prophage positions one might expect that the insertion of phages would be less directed, and more random. However, we found that the majority of phages inserted into two locations as seen by a bimodal density plot with a sharp separation between the two peaks. Prophage genome integration can either be a site-specific recombination event at so-called att sites or occurs in a non-directed manner by transposition into random sites (Ramisetty and Sudhakari 2019). Our data could indicate that the two areas in the genome contain most of the attachment sites for the majority of phages. A previously published analysis of *Salmonella* and *E. coli* genomes found a large number of distinctive phage integration loci; in the case of *Salmonella*, 24 loci were shared among 102 *Salmonella* phages, amounting to four phages statistically sharing one integration site. In case of *E. coli*, 58 distinctive integration loci were identified for 369 phages, with statistically 6.6 phages per site (Bobay et al. 2013). It might be reasonable to assume that *A. baumannii* contains similar numbers of attachment sites, although we have not analysed potential sites in the genomes we investigated. However, regardless of whether a phage inserts via one of the various attachment sites or randomly via transposition, only two “hot spots” were observed in our study. In addition to the explanation that attachment sites might be more frequent in these two sections of the bacterial genome, prophage insertion into segments crucial for e.g. over-all gene regulation, or into household genes, would be an evolutionary disadvantage and might therefore be less commonly found.

Bacteriophages are classified into 12 families (ICTV 2019). Pioneering work in taxonomy divided tailed phages into three classes based on the morphology of the phages; Myoviridae have long contractile tails, Siphoviridae have long non contractile tails, and Podoviridae have short tails. Recently, the International Committee on Taxonomy of Viruses (ICTV) expanded the order Caudovirales, describing tailed bacteriophages to include 6 additional families, i.e: Ackermannviridae, Autographiviridae, Chaseviridae, Demerecviridae, Drexlerviridae and Herelleviridae, taking additional characteristics into consideration such as genome sequence, gene content, protein homology and the host (Adriaenssens et al. 2020). When analysing the families of prophages in this population of *A. baumannii* strains, we observed a prevalence of Siphoviridae which constituted 57% of all identified prophages. Together with the next family of phages, Myoviridae, which consists 1/3 of all prophages, the two groups make up 90% of all prophages identified. Among the remaining 10%, the largest group belongs to Podoviridae. These ratios are very similar to the ones that have been reported in other studies, and also the ratio of the most commonly found phages in nature (Costa et al. 2018).

Interestingly, the ratio between ambiguous to active prophages in case of the ones that have been identified as Siphoviridae, 0.654, markedly differs from the ratio calculated for the prophages that belong to Myoviridae, 0.579. It is unlikely that Siphoviridae prophage sequences are less prone to mutations, as they should be occurring at random. However, a mechanism that would specifically “protect” Siphoviridae prophages might be the case if daughter cells, where mutations in the prophages occur, would have an evolutionary disadvantage. This would imply that prophages influence the host behaviour positively, which has previously been shown in some cases (Bondy-Denomy and Davidson 2014; Nanda et al. 2015; Loh et al. 2019). Could the most likely scenario be that the genomes of Myoviridae are possibly larger than those of the Siphoviridae, making them more prone to random mutations and deletions? However, the size comparisons of the genomic sequences of the prophages that we identified, does not support this possible explanation: The Siphoviridae sequences display distribution with one clear peak at around 20 kb followed by a fairly broad peak with a plateau and a shoulder towards larger genome sizes, ranging from 28 to 65 kb (Figure 6B). In contrast to this, the genomic size distributions of Myoviridae showed two peaks, one around 20 kb, the second around 42 kb (Figure 6C). The size estimations are corroborated by the findings of a previous study which estimated the genome sizes of Siphoviridae to be approximately 50 kb in average, with a broad distribution between 24–101 kb, while Myoviridae display smaller genomes or around 34 kb (Costa et al. 2018).

One question we could pose is why there is no broad distribution of prophage sizes, and why do we observe “peaks”? Can we conclude from this data that certain genome sizes are advantageous from an evolutionary standpoint? The arch-Myovirus might have had a certain size that proved to be sufficient for the successful persistence during the course of evolution. Only smaller increases or decreases of the genome allowed evolutionary success, and no gradual increase or decrease in genome size occurred. However, an evolutionary leap or jump might have happened at some point, which might have led to a major increase of genomic size, creating a new, second type of a Myovirus class which is represented in the second, larger peak. Starting from this size, again only smaller changes, decreases or increases with regards to the genomic size, may have occurred, preserving the sharp separation of each peak. It would be interesting to investigate if the smaller Myovirus display a prolate head as does T4. The increase volume of this geometry allows the packing of a larger genome, which might explain the possible separation in two sizes. To test this hypothesis, smaller Myoviruses should have non-prolate heads. Viral classification is a complex topic. Possibly the genome sizes might help to contribute to classifying of microbial viruses in the future.

While Prophage Hunter extracts prophage genomes from bacterial genomes, the platform is a web-based tool that also distinguishes between “active” and “ambiguous” prophage genomes (Song et al. 2019). The developers of Prophage Hunter have used experimental data and conducted induction experiments with mitomycin C, to validate the program’s output, showing its ability to hunt for “active”, inducible prophages. Yet, conclusions should not be hastily drawn to assume that all “active” prophages can definitively excise from the host genome to commence the bacteriophage lytic life cycle; false positives may still exist. In this regard, induction experiments should be conducted to confirm that “active” prophages can indeed produce active particles.

Prophages are an important source for acquiring new genetic information, including antibiotic resistance genes, for their bacterial host. Phage-mediated transfer of genes from donor to recipient cells, also called transduction, has been shown to be instrumental in the spread of AMR genes both *in vitro* and *in vivo* (Haaber et al. 2016). In our study, we also investigated antimicrobial resistance (AMR) genes that are embedded in prophage sequences. Previous studies on prophage diversity in *A. baumannii* had found AMR genes (also called: ARGs) in many prophages that were analysed (Costa et al. 2018, López-Leal et al. 2020). Yet despite this, it remains to be shown whether prophages confer antimicrobial resistance to its host in *A. baumannii*. Our observation illustrates that phages might represent important contributors in the process of AMR acquisition. However, it remains to be said that we found less than 5% of a publicly available, deposited sequence reads to contain the prophage-encoded ARGs we initially identified, arguing that phage transduction is possibly not the prevalent mode of AMR acquisition but is second to other mechanisms such as plasmid uptake via conjugation. Interestingly, despite viruses in general showing highly condensed genomes trying to pack essential information in small volumes, bacterial viruses seem to have co-evolved with their hosts and carry genes that are not directly required for the virus but are beneficial to the host and thus also to the prophage.

## Conclusion

Our study attempts to take an inventory of prophages in the important nosocomial pathogen *A. baumannii*. We have analysed the phylogeny of the prophages, their position in the host genome and characterised their lengths, identifying “successful” i.e. widely distributed phages, and the dominant families, Myoviridae and Siphoviridae.

Several prophage sequences contained genes coding for antimicrobial resistance genes. By mapping these genes in all deposited illumina *A. baummannii* sequence reads, we found that less than 5% of all available host sequences contain such prophage-embedded genes, indicating that transduction may not be the major contributor to the emergence of antimicrobial resistance.

## Supporting information

Supplementary sheet 1

Supplementary sheet 2

## Acknowledgments

We thank the National Science Foundation of China for supporting this work (NSFC 32011530116). BL was supported by the China Postdoctoral Science Foundation (530000-X91902). We would also like to thank the reviewers who made very valuable suggestions that helped to improve the work.

## Conflict of Interest

The authors declare that the research was conducted in the absence of any commercial or financial relationships that could be construed as a potential conflict of interest.

## Supplementary Material

The Supplementary Material for this article can be found online at:

## Supplementary Material

Supplemental Table S1: Metadata of all *A. baumannii* genomes used in this study.

Supplemental Table S2: Details of detected active and ambiguous prophage in all 177 *A. baumannii* strains analysed in this study.

Supplemental Figure S1: *In silico* detection and distribution of prophage sequences in *A. baumannii* strains. Unrooted core phylogenetic tree of 177 *A. baumannii* genomes from clinical isolates. Presence of prophage sequences are indicated by red squares. Green squares indicate the lack thereof. Please refer to the PDF of the figure and use the zoom function to identify names of strains and phages.

Supplemental Figure S2: Phylogeny of prophages found in *A. baumannii* genomes.

Supplemental Figure S3: Prevalence of prophages carrying AMR genes in *A. baumannii* strains.

## Notes

### Competing Interest Statement

The authors have declared no competing interest.

