## Supplementary figures and images for "A Biological Inventory of Prophages in *A. baumannii* Genomes Reveal Distinct Distributions in Classes, Length and Genomic Positions"

### Supplementary figure

Supplemental Figure S1

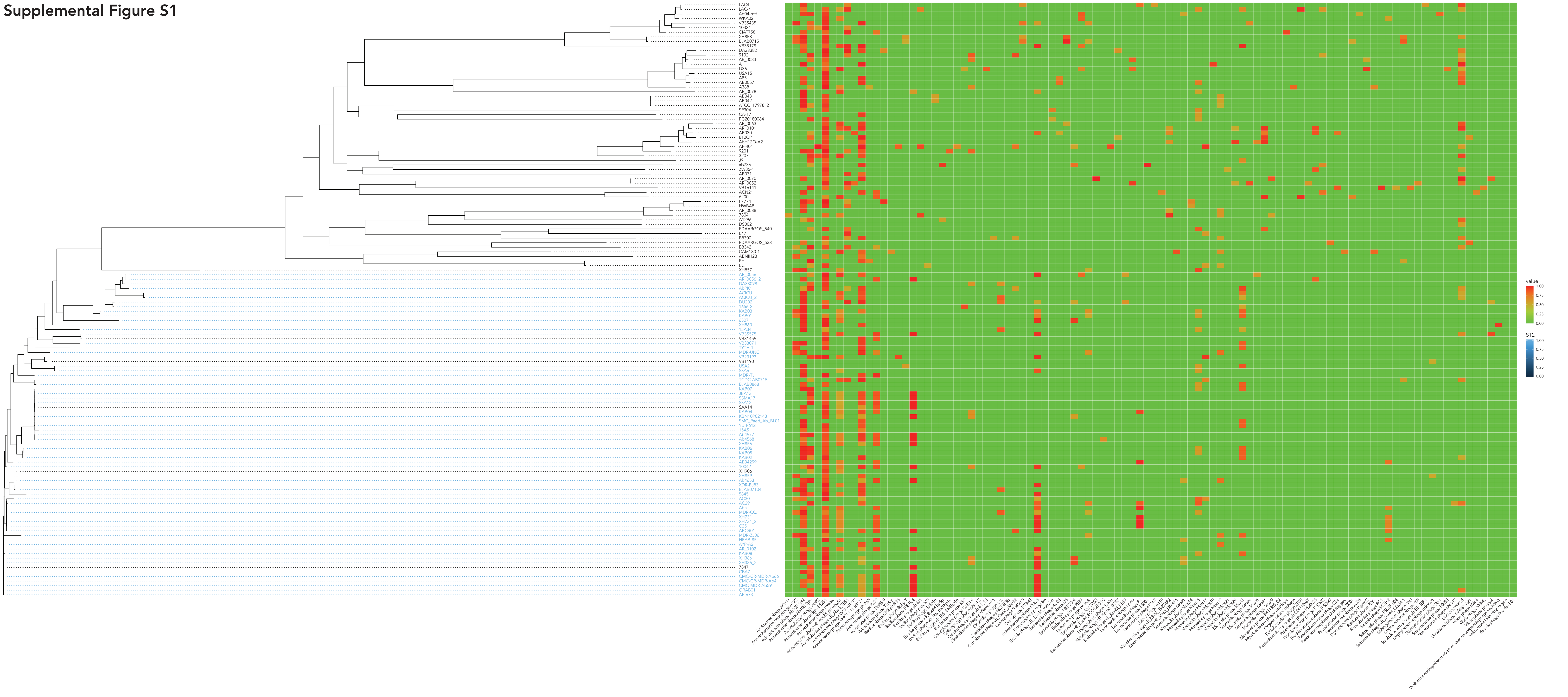

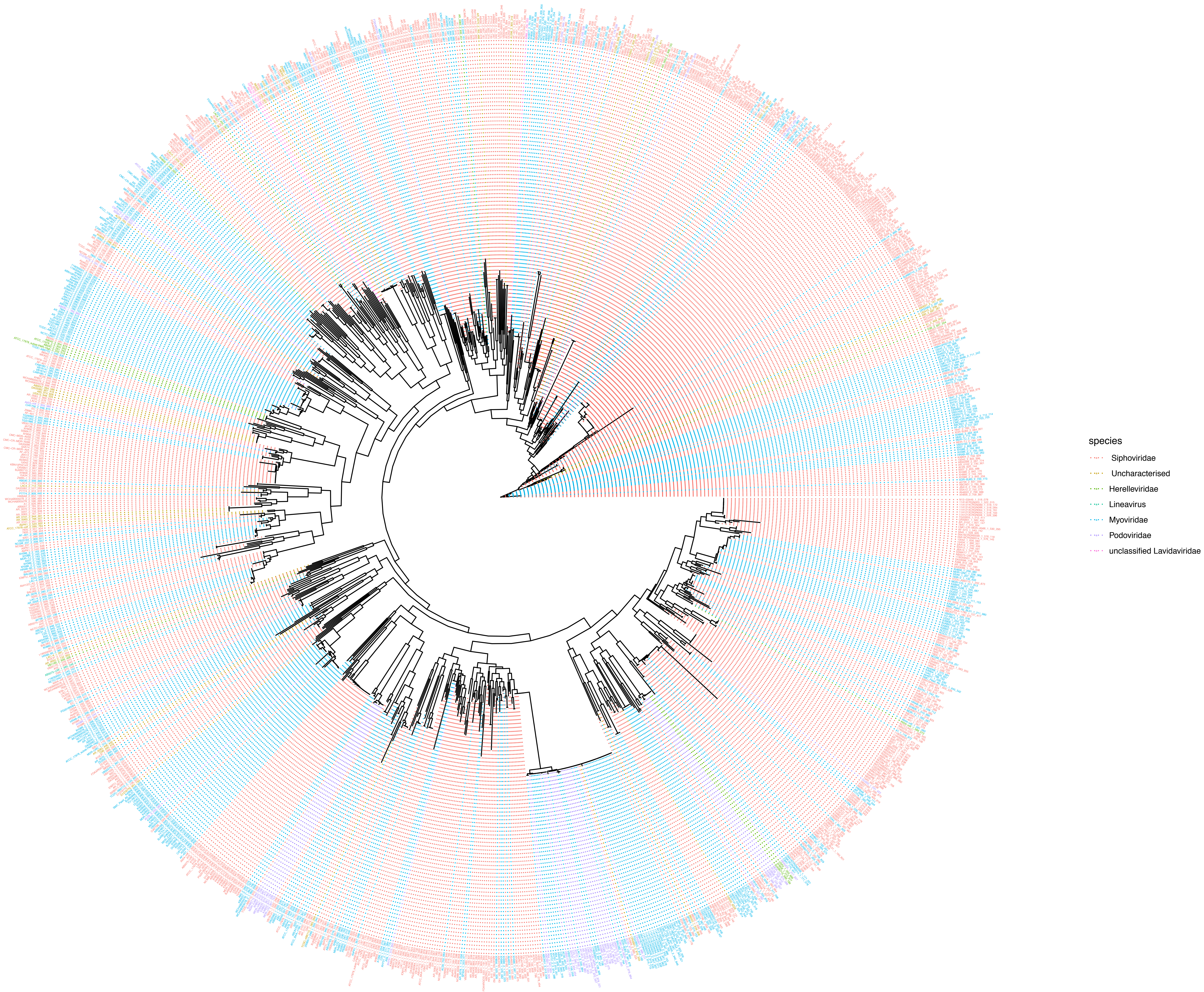

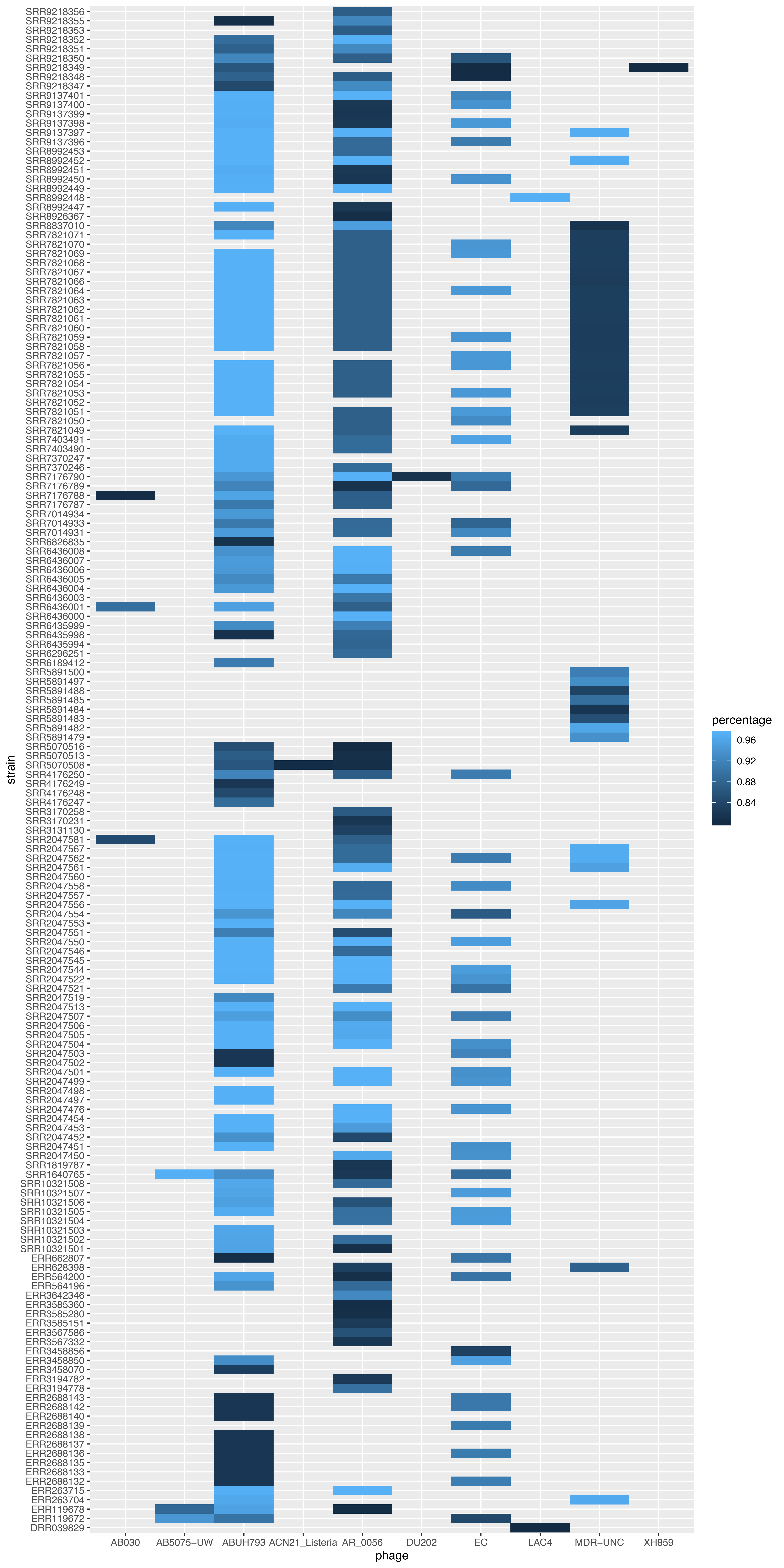
